# The impact of perturbation intensity schedule on improvements in reactive balance control in young adults: an experimental study

**DOI:** 10.1101/2025.02.05.636697

**Authors:** Piufat Wong, Isabel Yeh, Avril Mansfield, Christopher McCrum

## Abstract

**Background:** Reactive balance training (RBT) uses unanticipated perturbations to improve balance reactions and prevent falls. Intensity and predictability of perturbations may affect the generalizability of RBT, but their relative contributions remain unclear. This study aimed to compare the effects of three RBT schedules with differing intensity and predictability on participants’ balance recovery following untrained perturbations, and to determine perceived difficulty and challenge of the training schedules.

**Methods:** Participants were 36 healthy young adults (20-35 years). Participants were randomly assigned to one of three RBT intensity schedules (fixed high intensity, low-to-high intensity, and variable intensity). Training took place on a motion platform that delivered perturbations in varying directions (forward, backward, left, and right) and intensities across five trial blocks. Balance reactions (number of recovery steps) pre- and post-training, and electrodermal responses, and perceived difficulty during training were collected. Statistical analyses compared post-training outcomes between training groups, controlling for the baseline value.

**Results:** Participants took fewer steps and decreased the proportion of multi-step reactions pre- to post-training (p<0.0001), with no significant between-group differences. Perceived exertion decreased significantly across training blocks in the fixed-high group and increased significantly in the low-to-high group. Electrodermal responses declined across all groups between training blocks 1 and 3 (p=0.017).

**Conclusions:** Improvements in step reactions to untrained perturbations did not differ between training groups, highlighting the importance of multi-directional variability over specific intensity schedules for enhancing balance recovery.

## 1. INTRODUCTION

Reactive balance training (RBT; also referred to as perturbation-based balance training) uses unexpected balance perturbations to destabilise individuals and improve their balance (1, 2). Reactions to destabilising perturbations, which stabilise the position and velocity of the centre of mass relative to the base of support, are crucial for preventing a fall (3). Previous studies have found that RBT can improve reactive balance control and may prevent falls in daily life (1, 4, 5). However, these improvements in balance reactions can be specific to the tasks that have been trained, and there may be minimal transfer to untrained tasks (4). For instance, individuals who underwent gait-perturbation training on a treadmill did not demonstrate greater stability following a subsequent lean-and-lease perturbation when compared to the control group with no prior training (6). Since it is not feasible for RBT in the clinical setting to mimic all potentially fall-inducing events, it is essential to identify factors that can improve the generalizability of RBT and allow the balance recovery improvements made during training to be applied to unforeseen real-life balance challenges. These factors include perturbation characteristics, such as type, intensity, variability, unpredictability (6-9). Prior studies have suggested that generalizability is enhanced with RBT protocols using large magnitude and low predictability perturbations (7, 10). However, the relative contributions of predictability and intensity in facilitating generalizability have not been directly assessed. This has led to an absence of an evidence-based training protocol in clinical settings that effectively weighs these two factors for optimizing effects.

Our primary objective was to determine the effect of RBT protocols with different intensity schedules on participants’ balance recovery following perturbations in an untrained direction. Specifically, we compared the effects of three different intensity schedules (fixed high intensity, low-to-high intensity progression, and variable intensity) on responses to an untrained perturbation direction. We hypothesized that the variable intensity schedule is would lead to greater inter-task transfer, reflected by better performance of the untrained task, similar to how training multiple perturbation directions has been shown to lead to more direction-generalizable improvements in reactive balance control (11).

Higher intensity and more variable perturbations in training ma y be more challenging and difficult than progressively increasing perturbation intensity. Individuals’ perception of the difficulty of RBT can influence their training motivation and adherence, thus affecting their training improvements (12). Therefore, our secondary objective was to determine the effect of the different intensity schedules on participants’ difficulty or challenge of training. The variable intensity and fixed high intensity groups were expected to exhibit higher perceived difficulty and challenge of training.

## 2. METHODS

### 2.1 Study Design

This study was a multi-arm randomized experimental study with three parallel experimental training groups. The study was conducted at the Toronto Rehabilitation Institute, University Health Network, and was approved by the University Health Network Research Ethics Board (study ID: 18-5550).

### 2.2 Participants

Thirty-six healthy young adults aged 20-35 years were recruited from the community. Exclusion criteria were: past or current lower limb injuries or surgeries, diagnosed neurological or musculoskeletal conditions that restrict independent mobility, and previous involvement in perturbation platform studies. Participants provided written informed consent to participate in the study. This was a convenience sample based on the time period available for recruitment and measurement. A sensitivity power analysis indicates that this sample provides power (0.8) to detect effects of f>0.54.

### 2.3 Apparatus

We used a custom computer-controlled 6m x 3m motion platform, which was programmed to translate forward, backward, left, and right (Figure 1). Perturbation intensity was defined as the peak platform acceleration and categorized into five levels of increasing intensity. For forward perturbations (resulting in a backward loss of balance), levels 1 to 5 corresponded to peak accelerations of 1.67, 1.84, 2.17, 2.34 and 3 ms^-2^, respectively, and for the other three directions, levels 1 to 5 corresponded to peak accelerations of 2, 2.5, 2.75, 3.25 and 3.5 ms^-2^, respectively. Two perturbation waveforms were used: “standard” and “extended”; the standard waveform consisted of a 300 ms acceleration phase immediately followed by a 300 ms deceleration phase, whereas the extended waveform consisted of a 300 ms acceleration phase followed by a 600 ms deceleration phase. Inclusion of both waveforms helped to prevent participants from using the deceleration phase to regain stability (13).

**Figure 1.**
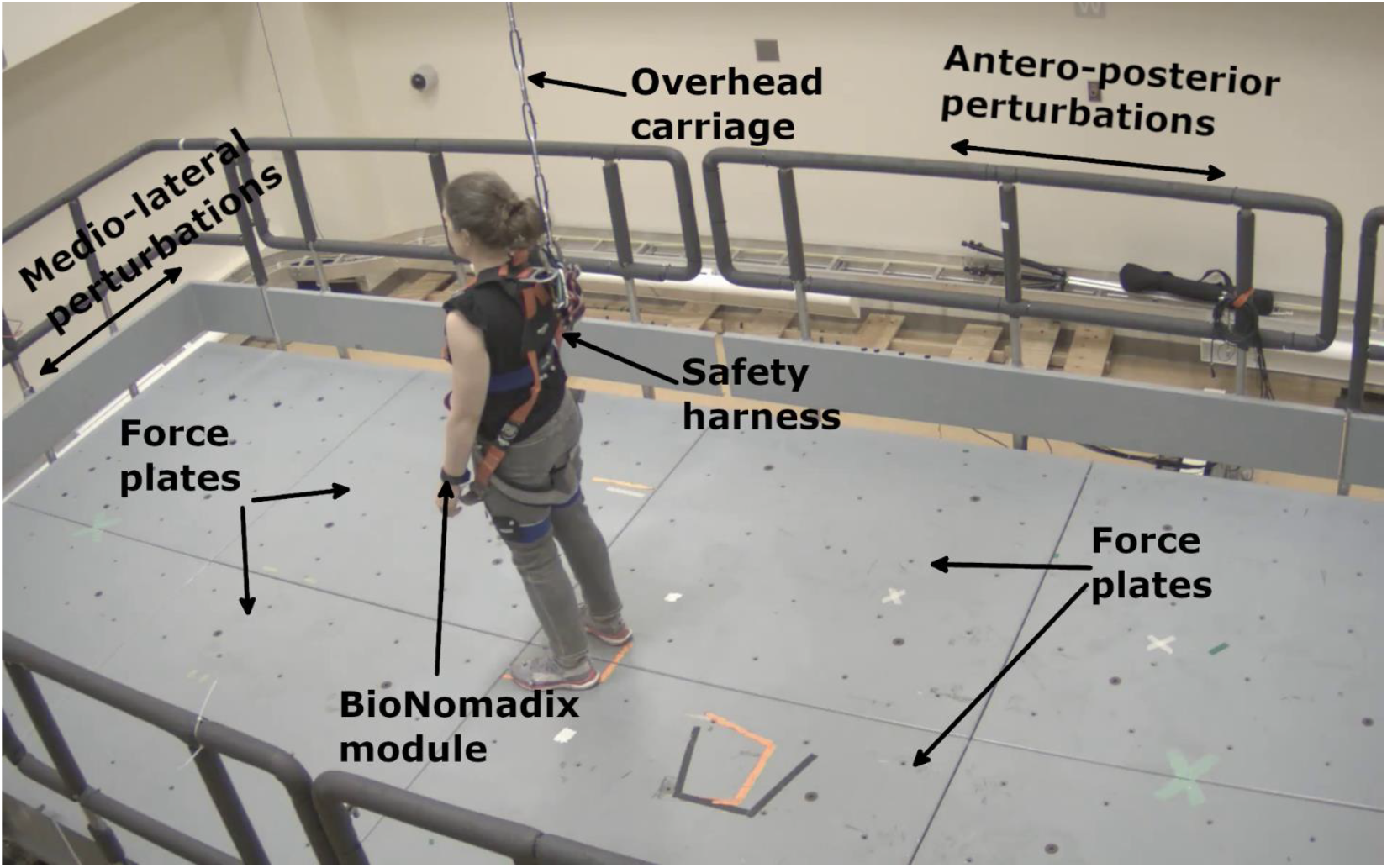
Experimental set-up. The participant stood in the standardized position with one foot on each of the two back force plates. The BioNomadix module was attached on the participant’s wrist to transmit the electrodermal activity signal to the receiver. A safety harness attached to an overhead carriage was worn to prevent falls. From the participant’s perspective, the directions of anterior-posterior and medial-lateral perturbations are identified in the figure with double-headed arrows.

Four 1.5m x 1.5m force plates (model BP11971197-2000, Advanced Mechanical Technology, Inc., Watertown, Massachusetts, USA) were installed on top of the motion platform to collect ground reaction forces and moments in three dimensions data at 1000Hz. Electrodermal activity was recorded wirelessly (BioNomadix module, BN-PPGED-T, BIOPAC Systems, Inc., Goleta, California, USA) at 1000Hz. Three-dimensional kinematic data were gathered with a 12-camera marker-based motion capture system (Vicon Vero 2.2, Vicon Motion Capture System Ltd., Oxford, UK) and an 8-camera markerless motion capture system (Theia 3D, Theia Markerless Inc., Kingston, Ontario, Canada) at 100Hz. Four additional video cameras were positioned around the room to record the session. An accelerometer (Series 7523A, Dynamic Transducers and Systems, Chatsworth, California, USA) placed on the platform recorded platform acceleration.

Participants wore a safety harness attached to an overhead carriage to prevent a fall to the ground. After fastening the harness, participants were instructed to attempt kneeling to ensure that their knees would not touch the ground if they fell. A load cell (SM500-10, Durham Instruments, Toronto, Ontario, Canada) in series with the harness line recorded any loading on the line. The harness offered no support unless participants started to fall.

### 2.4 Protocol

Each session lasted 1-2 hours, including obtaining informed consent, set up, data collection, and instrumentation removal. The following information was collected from participants at the beginning of the data collection session: age, biological sex, height, body mass, dominant hand and leg, and the Activities-specific Balance Confidence scale (14).

Eleven single reflective markers were placed over specific anatomical landmarks: 7^th^ cervical vertebra, greater trochanters, heels, on the shoes over the 2^nd^ and 5^th^ metatarsal phalangeal joints, and the tip of the big toe. Three marker clusters were securely attached to the participants’ thighs and upper chest with elastic straps. Furthermore, 7 markers were affixed on the platform. To collect electrodermal activity, two foam Ag/AgCl electrodes were placed on the palmar surface of the participants’ index and middle fingers on the non-dominant hand, which was first cleaned with alcohol wipes.

All data were recorded at least 5 seconds before each perturbation and continued for at least 5 seconds after the perturbation.

#### 2.4.1 Stance perturbations

Perturbations were delivered in five trial blocks: one pre-training test block, three training blocks (Figure 2), and one post-training test block (see further details below). Prior to each perturbation, participants stood in a standardized position (14 ° between the feet’s long axes and 17 cm between heel centres; 15) with one foot on each of the back force plates (Figure 1). They were asked to stand with their arms by their sides, facing forward; an ‘X’ was marked on the wall at participants’ approximate eye level for them to focus on during the tests. Before each perturbation, participants were asked if they were ready for the next trial. When participants indicated they were ready, they were told that the test was starting. Any time from 2-5 seconds following the warning, the platform moved; variability in the perturbation timing prevented participants from anticipating the perturbations. Participants were instructed to do whatever was necessary to recover their balance after the perturbation, and to hold their final position when they felt stable. The trial ended approximately 5 s after the perturbation, at which point participants were asked to rate the perceived difficulty of the perturbation using the OMNI Perceived Exertion Scale (16), which ranges from 0 (extremely easy) to 10 (extremely hard). Participants were then instructed to return to the starting position and get ready for the next trial.

**Figure 2.**
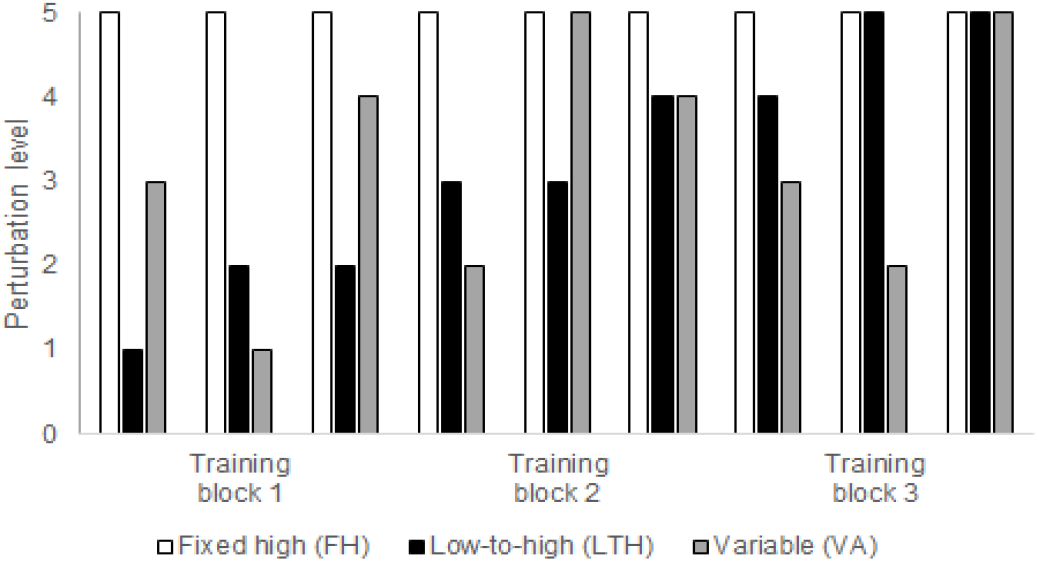
Training protocol. Participants in the Fixed high (FH) group experienced perturbations at Level 5 for all Training Blocks. Participants in the Low-to-high (LH) group experienced perturbations starting at Level 1 and increasing to Level 5. Participants in the Variable (VA) group experienced the same perturbation intensities as participants in the LH group, but in a random order across the three Trial Blocks.

##### 2.4.1.1 Pre- and post-training test

Pre- and post-training test blocks consisted of 10 perturbations: four backward perturbations with peak acceleration of 3.5 m/s^2^, and six perturbations to the left, right, or forward (2 per direction) with peak acceleration of 1.0 m/s^2^. The backward perturbation was used as an untrained task to assess inter-task transfer across the three groups; very small magnitude perturbations in other directions were included in the test block to reduce the predictability of the perturbations, while providing minimal training stimulus due to the low intensity.

After the post-training test, participants completed a short questionnaire with a 7-point Likert scale ranging from “Strongly agree” to “Strongly disagree” to measure their overall perceptions regarding the stressfulness, difficulty, and predictability of the perturbations experienced during the session (Appendix 1).

##### 2.4.1.2 Training

During the training blocks, participants experienced 27 platform movement perturbations in three directions (forward, left and right). Perturbations were presented in three blocks of nine perturbations (Figure 2), with scheduled rest breaks in between blocks; participants could also request a rest break at any time. The perturbation direction and acceleration waveform for all three groups followed the same predetermined sequence. The only difference between the groups was their intensity schedules: the fixed high intensity (FH) group experienced level 5 intensity for all perturbations; the low to high intensity (LH) group experienced increasing intensity levels from 1 to 5; and the variable intensity (VA) group experienced intensity levels from 1 to 5 in a varied order (Figure 2). Importantly, the number of perturbations at each intensity was matched between the VA and LH groups to ensure that we could determine that any differences between these groups were schedule-related and not due to differences in average perturbation intensity. Participants were randomly allocated to one of the three training groups using sealed envelopes.

### 2.5 Data processing

Marker-based kinematic data were labelled using Vicon Nexus (version 2.16, Vicon Motion Capture Systems Ltd., Oxford, UK). The Theia 3D proprietary algorithm used video footage of participants to scale a kinematic model template to their bodies based on the length of body segments, which was further adjusted by participant height in Visual 3D (version 2021.09.1, C-Motion, Germantown, Maryland, USA). Whole-body centre of mass (CoM) was calculated in Visual 3D using the default Theia model (17). Unfiltered marker and CoM data were exported for further processing using a custom routine implemented in MATLAB (The Math Works Inc., version 2022a, Natick, Massachusetts, USA).

Force plate and accelerometer data were low pass filtered at 20Hz using a 2^nd^ order zero phase-lag Butterworth filter. Electrodermal data were low-pass filtered at 2 Hz with a 2^nd^ order Butterworth filter. Kinematic marker and CoM data were low pass filtered at 6Hz with a 2^nd^ order dual-pass Butterworth filter. Kinematic data were synchronized with all other data using the platform acceleration signal (i.e., from acceleration of floor markers and the accelerometer). Perturbation onset time was the time when the platform acceleration exceeded 0.1ms^-2^ (18).

The primary outcome was the effectiveness of participants’ balance reactions following the perturbation in the untrained direction in both the pre-and post-training tests, as assessed by the number of recovery steps required to recover balance and the proportion of multi-step reactions. Additional explanatory outcomes were: foot-off time, swing time, step length, step width, centre of mass (CoM) displacement, margin of stability in anteroposterior (AP-MoS) and mediolateral (ML-MoS) directions (19), electrodermal responses (EDRs), and OMNI perceived exertion scale scores.

Recovery steps are the rapidly executed steps taken to regain balance following a large balance perturbation (20). Video footage was reviewed to count the number of recovery steps taken. A multi-step reaction was defined as taking two or more steps to recover balance. Foot-off time was defined as the time between perturbation onset and when vertical force recorded by the force plate under the stepping limb was less than 1% of participants’ body weight. Foot contact time was defined as the time after foot-off when the vertical force recorded by the force plate under the stepping limb was more than 1% of participants’ body weight. Vertical forces were visually inspected and foot-off or foot-contact times were adjusted if necessary (e.g., in the case of noise in the signal). Swing time was the difference between foot-off and foot-contact time. Step length and width for the first recovery step were the anteroposterior and mediolateral, respectively, difference in the position of the heel markers at foot contact. Motion platform displacement (from the floor markers) was subtracted from the CoM. CoM displacement was the difference in the anteroposterior position of CoM at the time of foot contact and perturbation onset. The extrapolated CoM (XcoM) in the anteroposterior and mediolateral directions were calculated as follows (19):

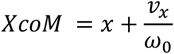

where *x* is the vertical projection of the CoM (m) on the ground, *v*_*x*_ is the CoM velocity (m/s), 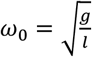 is the eigenfrequency of the pendulum, *g* is the gravitational constant (9.81ms^-2^), and *l* is the effective pendulum length (m) for an inverted pendulum model (1.24 times (21) or 1.34 times (22) participants’ trochanteric height for antero-posterior and medio-lateral XcoM, respectively (19, 23). Centre of pressure (CoP) was calculated under the stepping limb from the forces and moments recorded by the force plates. Margin of stability was then calculated as the difference between the CoP and XcoM (19, 24); margin of stability was calculated at 25% loading of the stepping limb due to low vertical forces leading to error in calculating the CoP location at foot contact. For a backward perturbation with a forward recovery step, a positive AP-MoS indicated that the CoP was in front of the XcoM. The sign of the ML-MoS was adjusted for left and right steps such that a positive ML-CoP indicated that the CoP was more lateral than the XcoM. The baseline electrodermal level was the electrodermal activity 1 s prior to perturbation onset. EDR was the peak in the electrod ermal signal up to 10 s post-perturbation, minus the baseline electrodermal level.

### 2.5 Data analysis

Participant characteristics were compared between groups using one-way analysis of variance (ANOVA; continuous variables, except ABC), Kruskal-Wallis test (ABC score), or Fisher’s exact test (categorical variables). To determine participants’ arousal following the training perturbations (which relates to challenge of responding to the perturbation (25)), and perceptions of difficulty of the training, two-way repeated measures ANOVA was used to compare EDRs and OMNI perceived exertion scores between groups during training; factors in the ANOVA were group (three levels), trial block (three levels: block 1, 2 and 3; see Figure 2), and the group-by-trial block interaction. Kruskal-Wallis test was used to compare perceptions of perturbations scores between groups. One-way analysis of covariance (ANCOVA) was used to compare primary and secondary outcomes post-training between the three groups, with the baseline value as a covariate. Outcomes were rank transformed prior to ANCOVA to allow for non-parametric analysis (26). Wilcoxon signed rank test was used to compare outcomes pre- and post-training across all groups combined. To probe short-term within-block adaptation versus inter-task transfer due to training, we also conducted a two-way repeated measures ANOVA comparing average number of steps and proportion of multi-step reactions between the last two pre-training trials and the first two post-training trials; factors in the ANOVA were group (three levels), trial (4 levels: 3^rd^ and 4^th^ trial pre-training, and 1^st^ and 2^nd^ trial post-training) and the group-by-trial interaction effect. The criterion significance level was set at p<0.05 for all analyses. All statistical analysis was conducted using Stata (StataNow/SE 18.5, StataCorp LLC, College Station, Texas, USA).

## 3. RESULTS

### 3.1 Participant characteristics

The groups did not differ in baseline characteristics: age, weight, height, proportion of sex, dominant hand and foot, and ABC score (Table 1).

**Table 1:** Participant characteristics. Values presented are means with standard deviations in parentheses for continuous variables, or counts for categorical variables. The p-value is for the one-way ANOVA (continuous variables, except ABC), Kruskal-Wallis test (ABC score), or Fisher’s exact test (categorical variables), comparing the three groups.

|  | Fixed high | Low-high | Variable | p-value |
| --- | --- | --- | --- | --- |
| Age (years) | 24.7 (3.2) | 27.3 (5.2) | 28.4 (4.4) | 0.11 |
| Sex (number female) | 8 | 6 | 5 | 0.49 |
| Height (m) | 1.67 (0.07) | 1.67 (0.11) | 1.68 (0.06) | 0.97 |
| Mass (kg) | 69.8 (14.3) | 65.1 (13.8) | 64.8 (8.6) | 0.56 |
| Dominant hand (number right) | 12 | 9 | 12 | 0.092 |
| Dominant foot (number right) | 12 | 8 | 11 | 0.10 |
| ABC score (%) | 88.2 (10.1) | 90.7 (7.4) | 93.4 (6.9) | 0.32 |

ABC: Activities-specific Balance Confidence scale.

### 3.2 Missing data and protocol deviations

Due to human error, one participant in the FH group only completed 3 backward perturbations in the pre-training test block. Therefore, analysis of number of steps, frequency of multi-step reactions, foot-off time, swing time, step length, step width, and OMNI scores is based on 287 trials. EDRs were not observed in 4 trials for two participants (both FH group) and due to technical error electrodermal data were not collected for one participant (LH group); therefore, analysis of EDRs is based on 275 trials across 35 participants. Due to technical error, markerless motion capture data could not be processed for one trial (post-training, FH group); therefore, analysis of CoM and MoS is based on 286 trials. Two participants (one FH group, one LH group) declined to answer one question regarding perceptions of perturbations (“I enjoyed taking part in the training”); therefore, analysis of this variable is limited to 34 participants. Due to human error, five participants in the LH group and two participants in the VA group did not complete training per protocol. These participants experienced left and right perturbations at 2.0 m/s^2^ when the protocol dictated perturbations at 3.0 m/s^2^ and experienced forward perturbations at 3.0 m/s^2^ when the protocol dictated perturbations at 2.0 m/s^2^. This error affected 6 perturbations in Trial Block 2 for the LH group, and 3 perturbations in Trial Block 1 and 3 perturbations in Trial Block 3 for the VA group. To confirm the results were unaffected by this protocol error, analyses were repeated with these participants excluded (see Appendix 2).

### 3.2 Arousal and perceptions of perturbations

There was a significant group-by-training block interaction effect for OMNI perceived exertion scores (F_4,33_=12.35, p<0.0001; Figure 3). OMNI perceived exertion scores significantly declined between Training Block 1 and 2, and between Training Block 1 and 3 for the FH group, whereas they significantly increased between Training Blocks 1 and 2, and between Training Blocks 2 and 3 for the LH group. There were no significant differences in OMNI perceived exertion scores across training blocks for the VA group. There were no significant differences in OMNI perceived exertion scale scores between groups across any of the training blocks.

**Figure 3:**
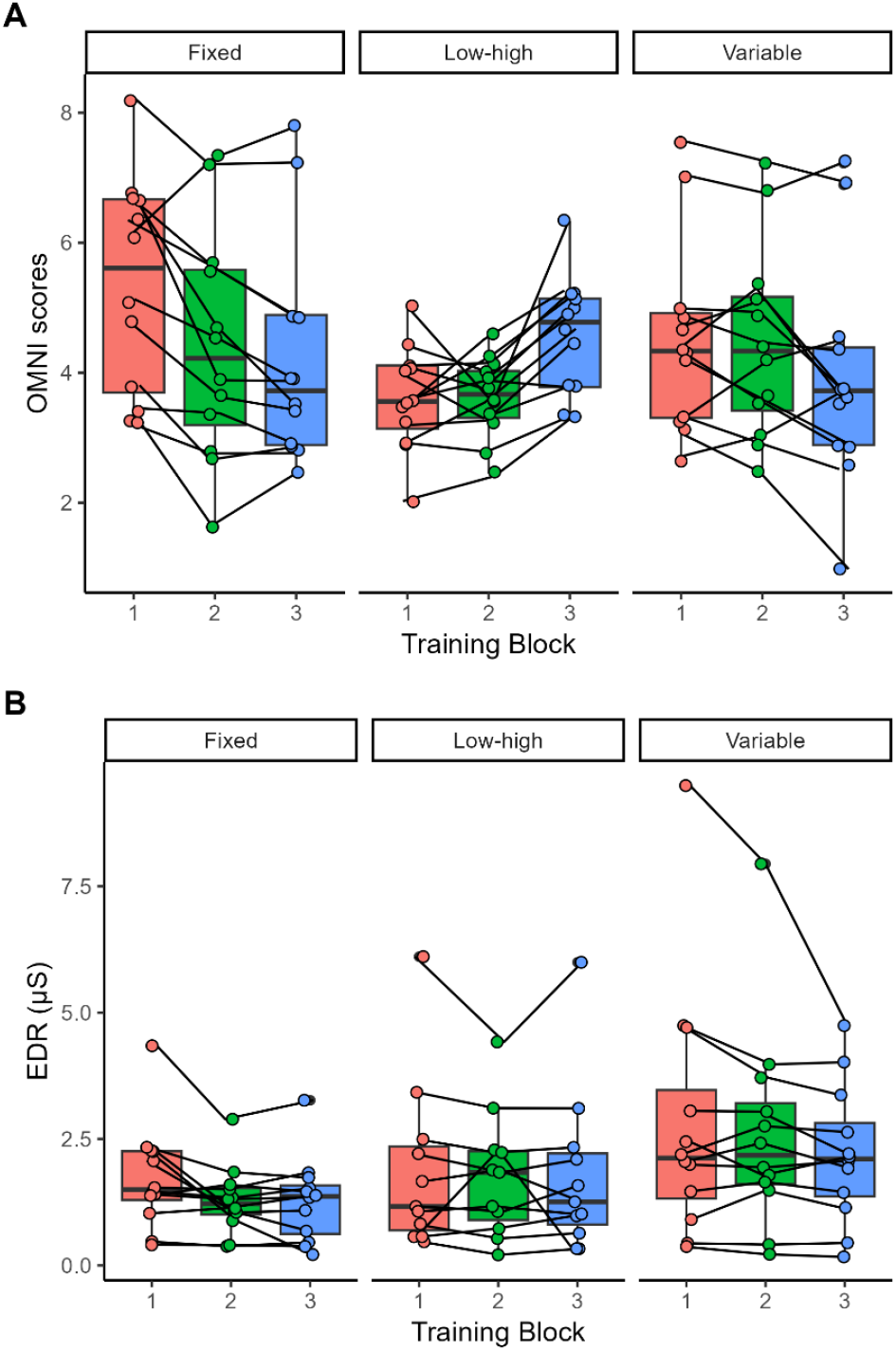
Arousal and perception of perturbations during training. OMNI perceived exertion scores (Panel A) and electrodermal responses (EDR, Panel B) for each group during the training blocks. Values plotted are individual data points for each participant, with line connecting data points for each participant. Box-plots show the group medians, and first and third quartiles.

There was no significant group-by-training block interaction effect (F_4,32_=1.06, p=0.38) and no significant main effect of group for EDR (F_2,32_=1.72, p=0.20; Figure 3). There was a significant effect of training block for EDR (F_2,32_=4.35, p=0.017). Post-hoc testing revealed that EDR magnitude declined across all groups between Training Blocks 1 and 3.

There were no significant differences between groups in perceptions of the perturbations at the end of the training session (Figure 4).

**Figure 4:**
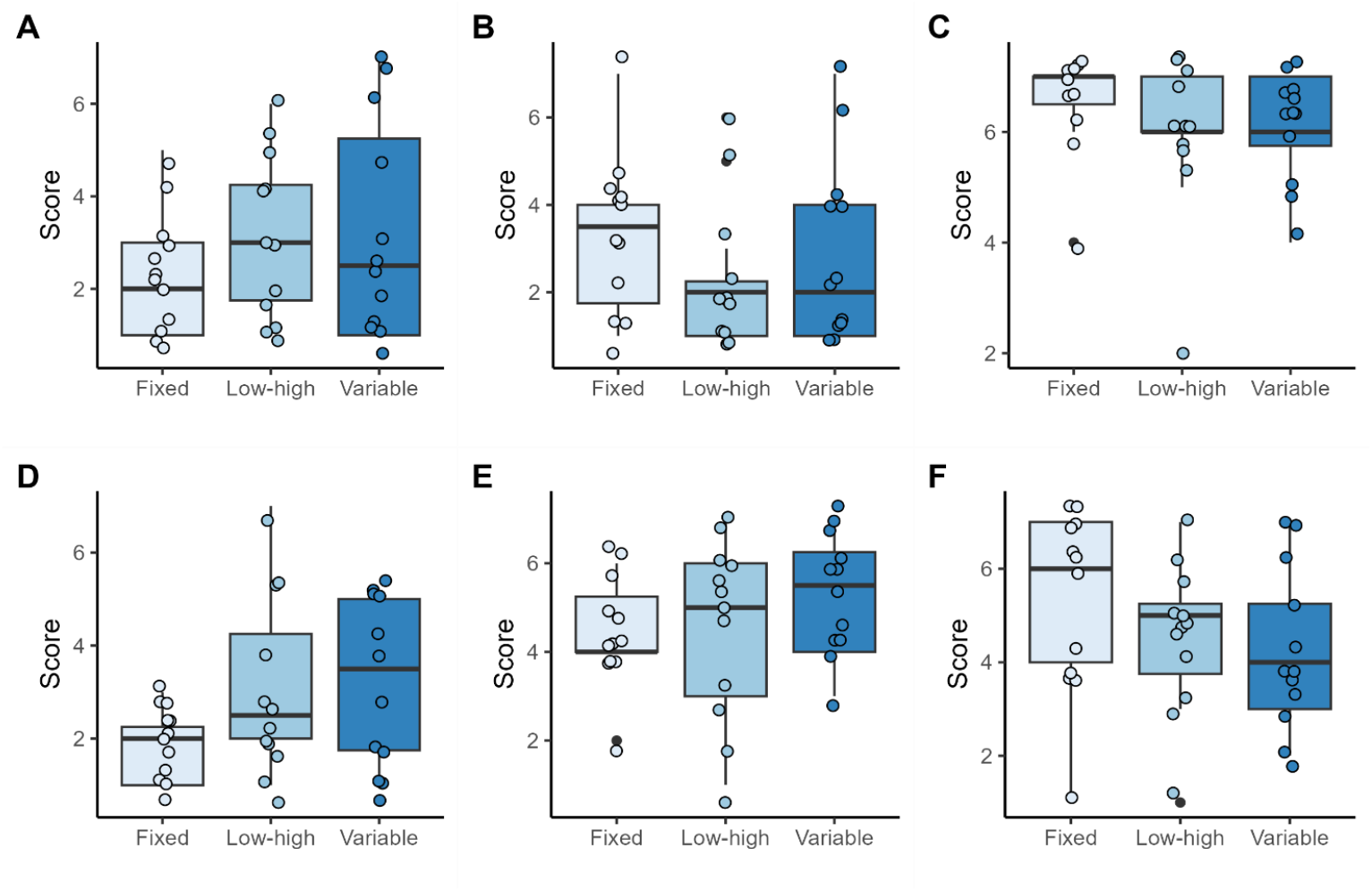
Perception of perturbations. Scores for ‘Stressful’ (Panel A; χ^2^=1.05, p=0.60), ‘Easy to predict the next task’ (Panel B; χ^2^=1.38, p=0.50), ‘Enjoyment’ (Panel C; χ^2^=2.40, p=0.30), ‘Difficulty’ (Panel D; χ^2^=3.62, p=0.16), ‘Tasks were expected’ (Panel E; χ^2^=1.93, p=0.38), and ‘Did not find training challenging’ (Panel F; χ^2^=2.44, p=0.29) for each group post-training. Values plotted are individual data points for each participant. Box-plots show the group medians, and first and third quartiles. Scores range from 1-7 indicating agreement with the statement, with lower scores indicating stronger disagreement, and higher scores indicating stronger agreement.

### 3.3 Primary outcome: step reactions

From ANCOVA, there was no significant between-group difference in number of steps (F_2,32_=1.28, p=0.29) or proportion of multi-step reactions (F_2,32_=0.14, p=0.87; Figure 5) post-training. Wilcoxon signed rank tests found that, overall, participants took fewer steps (Z=4.53, p<0.0001) and fewer multi-step reactions (Z=4.53, p<0.0001) post-training compared to pre-training.

**Figure 5:**
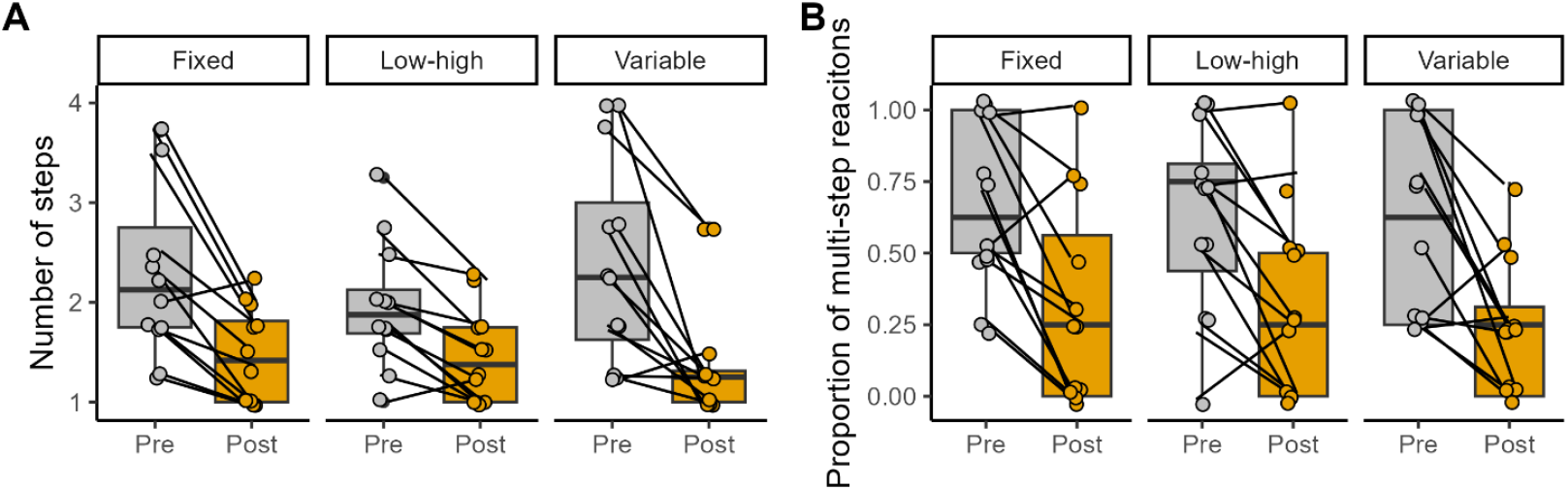
Primary outcomes (step reactions). Average number of steps (Panel A) and proportion of multi-step reactions (Panel B) for each group pre- and post-training. Values plotted are individual data points for each participant, with lines joining the pre- and post-training values for each participant. Box-plots show the group medians, and first and third quartiles.

When examining outcomes with the testing trial blocks, two way repeated measures ANOVAs found no statistically-significant group-by-trial interactions for average number of steps (F_6,33_=0.89, p=0.51) or number of multi-step reactions (F_6,33_=0.43, p=0.85; Figure 6), but there were significant main effects of trial order for both outcomes (F_3,33_>3.74, p<0.014). Post-hoc testing revealed that average number of steps significantly declined between the 3^rd^ pre-training trial and the 2^nd^ post-training trial, and proportion of multi-step reactions was significantly lower for the 2^nd^ post-training trial compared to the 1^st^ post-training trial, and the 3^rd^ and 4^th^ pre-training trials. Average number of steps and proportion of multi-step reactions did not differ between the 3^rd^ and 4^th^ pre-training trials, or between the 4^th^ pre-training trial and 1^st^ post-training trial.

**Figure 6:**
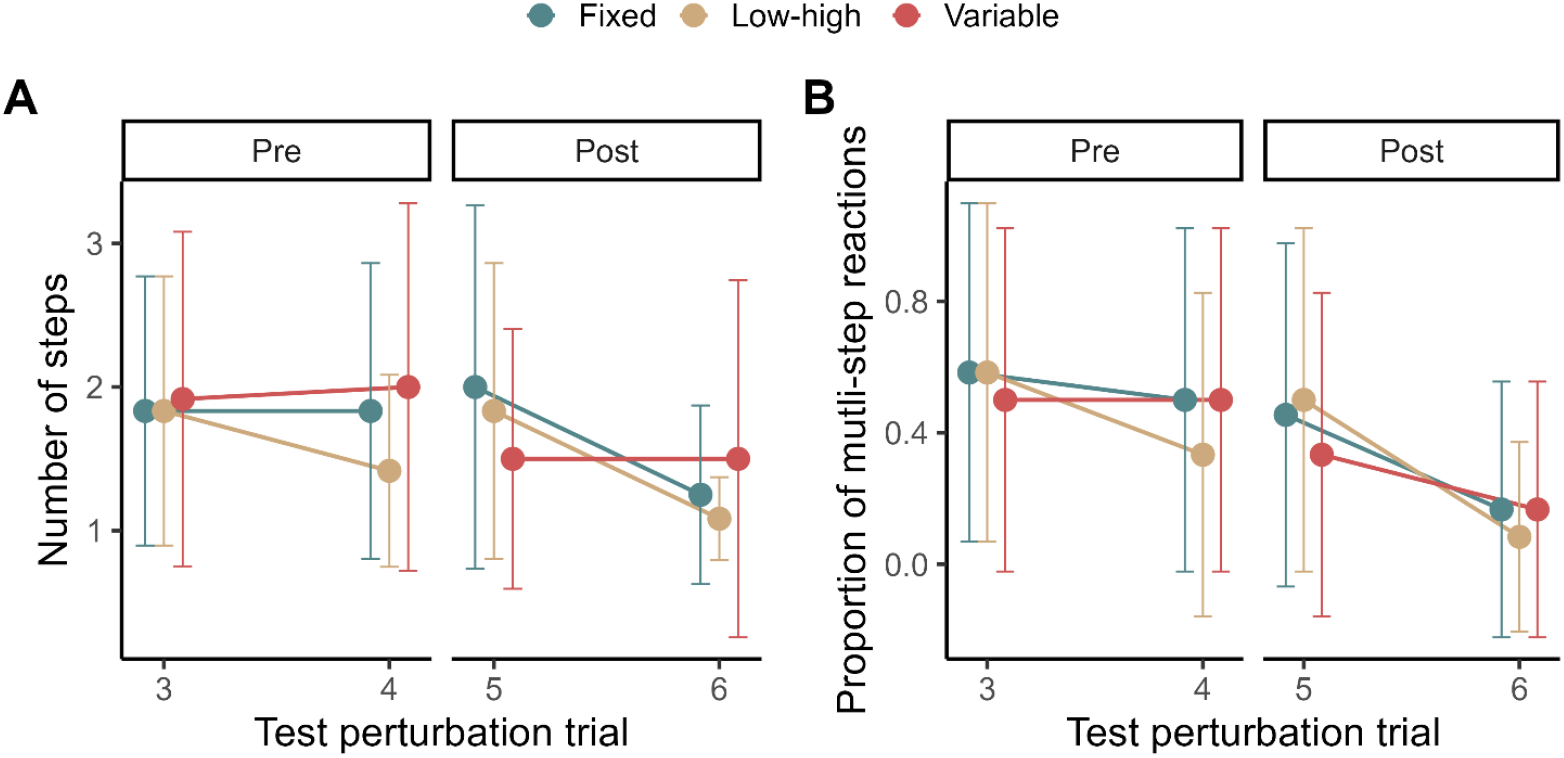
Step reactions for individual trials. Number of steps (Panel A) and proportion of multi-step reactions (Panel B) for the last 2 pre-training perturbations (trials 3 & 4), and the first 2 post-training trials (5 & 6). Values plotted are means with standard deviation error bars.

### 3.4 Secondary outcomes

From ANCOVA, there were no significant between-group differences in any CoM-derived variables post-training (CoM displacement: F_2,32_=0.80, p=0.46; AP-MoS: F_2,32_=0.28, p=0.76; ML-MoS: F_2,32_=1.06, p=0.36; Figure 7). There was a significant increase in ML-MoS pre- to post-training across all groups (Z=3.45, p=0.0003), but no significant change pre- to post-training for CoM displacement (Z=1.30, p=0.20) or AP-MoS (Z=0.96; p=0.35).

**Figure 7:**
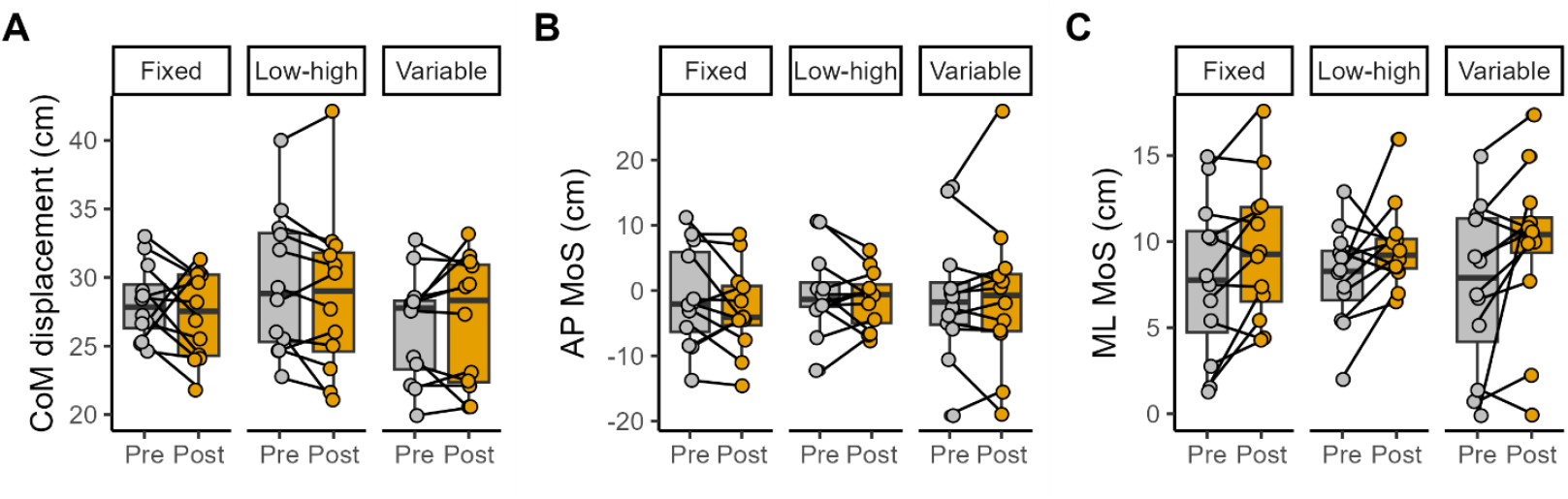
Secondary outcomes (CoM-derived variables). CoM displacement (Panel A), AP MoS (Panel B), and ML MoS (Panel C) for each group pre- and post-training. Values plotted are individual data points for each participant, with lines joining the pre- and post-training values for each participant. Box-plots show the group medians, and first and third quartiles.

There were no significant differences between groups in any spatio-temporal variables post-training (foot-off time: F_2,32_=1.01, p=0.37; swing time: F_2,32_=0.04, p=0.96; step length: F_2,32_=0.20, p=0.82; step width: F_2,32_=0.30, p=0.74; Figure 8). Likewise, there were no significant changes pre-to post-training for any spatio-temporal variables (foot-off time: Z=1.28, p=0.20; swing time: Z=1.45, p=0.15; step length: Z=0.75, p=0.46; step width: Z=1.87, p=0.062).

**Figure 8:**
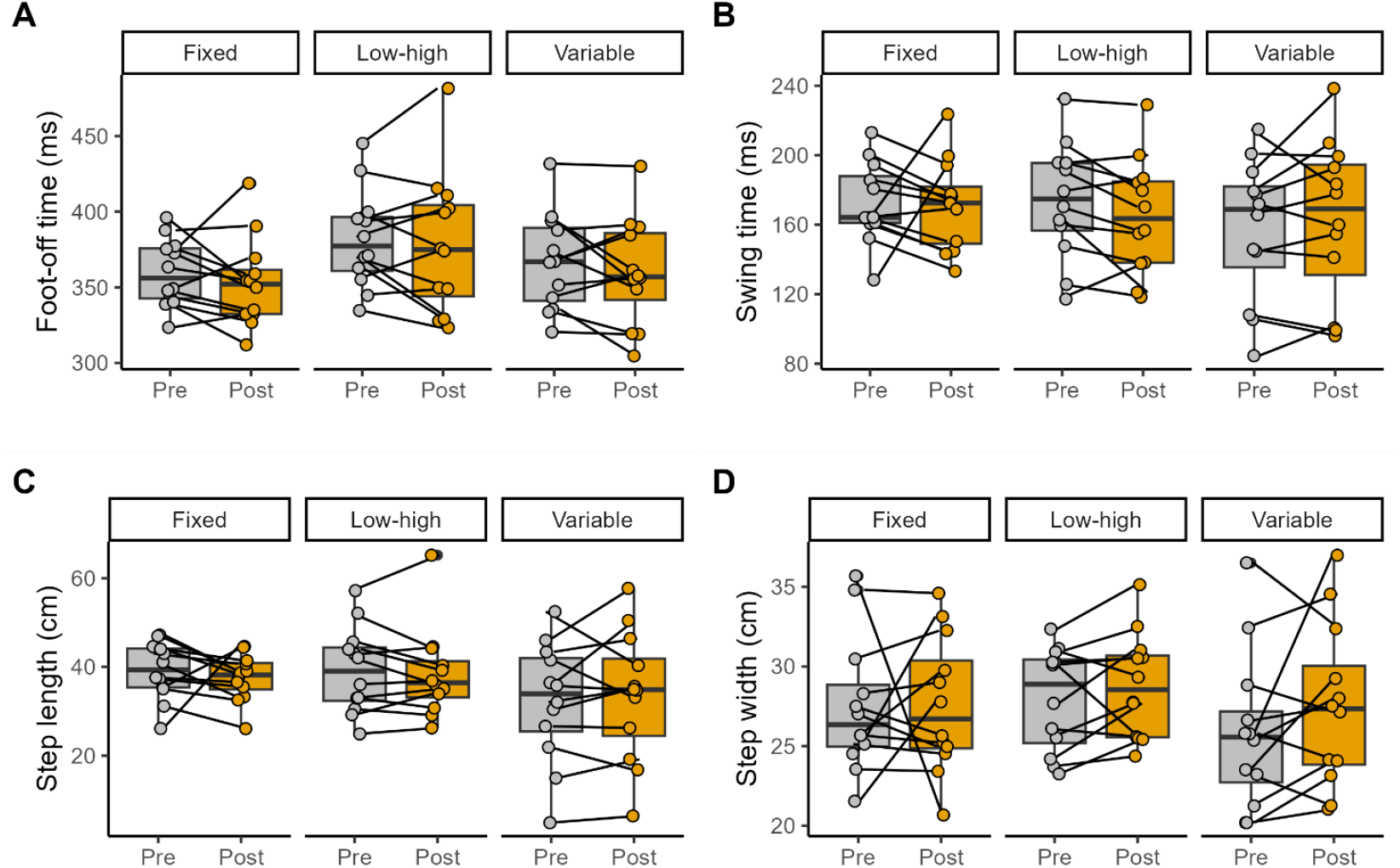
Secondary outcomes (spatio-temporal variables). Foot-off time (Panel A), swing time (Panel B), step length (Panel C), and step width (Panel D) for each group pre- and post-training. Values plotted are individual data points for each participant, with lines joining the pre- and post-training values for each participant. Box-plots show the group medians, and first and third quartiles.

### 3.5 Results for participants with protocol deviations excluded

When analysis was repeated with participants with protocol deviations excluded, the findings were generally the same as for the full sample (Appendix 2). With the reduced sample, the increase in OMNI perceived exertion scores across Training Blocks for the LH group and the decline in proportion of multi-step reactions between the 3^rd^ and 4^th^ pre-training trials were not statistically significant.

## 4. DISCUSSION

This study aimed to investigate the effects of RBT protocols with different intensity schedules on participants’ balance performance following a perturbation in a new direction. Our secondary aim was to determine the challenge of the different RBT protocols, and perceptions of training difficulty. Contrary to our hypotheses, we found that all groups improved their reactive stepping and reduced physiological arousal post-training, with no effect of training schedule on outcomes.

All groups improved in their step responses. Contrary to the hypothesis, the different intensity schedules did not appear to have a measurable influence on the primary outcomes. Previous work has shown that higher-magnitude and/or less predictable training schedules induce better motor learning, but only for same-direction perturbations that differ in condition (7, 10) or magnitude (27), or for same-plane perturbations that differ in direction (28). There is little direct evidence of the effects of varying intensity training schedules that combine perturbations in both the anterior-posterior and medial-lateral directions. Perturbations in different directions demand different balance responses. The neural pathways needed to recruit the muscle groups associated with a forward perturbation, for example, differ from those for a leftward perturbation (2, 29). Therefore, it is possible variability and unpredictability in direction of the training perturbations had a larger effect on adaptation than intensity schedules. This suggests that variable direction of training perturbations may be more important than variable intensity for improving balance reactions.

The average number of steps decreased, suggesting that participants became more efficient in their balance reactions after training. These results confirm the ability of healthy adults to improve balance reactions with a short period of practice (28, 30, 31). Most participants took no more than one recovery step post-training, displaying minimal room for further improvement. Our results also indicate that training in left, right, and forward directions transferred to improved quality of reactions in the backward direction. This contrasts previous work by others (32), where effects of training with perturbations in the anterior-posterior direction did not transfer to untrained perturbations in the medial-lateral direction. The lack of transfer shown here suggests that the movement patterns or neuromuscular strategies required for left and right perturbations were not sufficiently similar to those required for forward and backward perturbations. This idea aligns with the recommendations of a previous review, which suggests that multi-directional perturbations in training are necessary to generalize improvements to multiple contexts (33). In motor learning research, the relationship between variability and generalizability is well established, with evidence that variability in practice promotes greater, more errorless transfer of skills to novel tasks compared to constant practice (34). Therefore, it seems that greater variability of perturbation directions is beneficial for improving generalization to untrained perturbations.

The increase in ML MoS from pre- to post-training indicates a greater distance between CoP and XcoM, and therefore improved stability in the ML direction (19). Interestingly, AP stability was not shown to improve, despite the test perturbations occurring in the AP direction. Similar results are sparse in the current literature, but it is plausible that this improvement in the ML stability, and lack thereof in the AP direction, may simply be because there were twice as many ML training perturbations compared to AP. This increased exposure would have allowed for more training of ML CoM and control of foot positioning. Large AP perturbations that require reactive steps can disrupt ML stability as the reaction must be executed quickly, without sufficient time for stabilizing anticipatory postural adjustments (35, 36). Consistent with previous studies in older adults, our results suggest that young adults can learn to control this lateral instability with practice responding to balance perturbations (4). We did not observe a significant increase in step width following training; therefore, improved ML stability was primarily achieved by slowing lateral motion of the COM, rather than increasing the size of the base of support.

Balance perturbations evoke EDRs, which are phasic increases in electrodermal activity that occur 2-5 seconds post-perturbation (37). While the role of these EDRs in balance reactions is unclear (25), they are thought to represent increased physiological arousal following the perturbation that relates to the complexity or challenge of the response (38). There was no significant between-group difference in EDRs, suggesting that the challenge of training schedules may not have been markedly different between groups. This finding is supported by the lack of a between-group difference in perceived difficulty of training based on OMNI perceived exertion scores. EDRs declined during training for all three groups. This finding is consistent with a prior study showing reduced electrodermal activity as participants practiced and improved maintaining balance on a moving platform with unpredictable continuous balance perturbations (39). Therefore, reduced EDRs in the current study suggest reduced difficulty of the responding to the perturbations as participants’ adapted with practice.

There were no differences between groups in the perceived “stressfulness” of the perturbations. These results conflict with those from Okubo et al. (9), where more unpredictable perturbation training during walking trials was correlated with greater anxiety. Training sessions in the study by Okubo et al. (9) consisted of six disturbances, with each session progressively increasing in level of unpredictability of perturbation characteristics. Participants in the current study experienced a greater number of perturbations that remained stable in their levels of unpredictability throughout training, allowing for more practice and exposure to the demands of the task. It should be noted that Okubo et al. (9) a) did not examine the influence of perturbation intensity on anxiety, so the relative effects of perturbation intensity and number cannot be compared directly, and b) applied their perturbations during walking, which involves greater velocities and perhaps a higher perceived threat of a fall. Prior research shows that repeated exposure to postural threats reduces anxiety responses over time (40, 41). As individuals gain more experience successfully managing balance disturbances, their perceived risk of falling diminishes. This principle has been used extensively in exposure-based therapies, in which individuals undergo repeated threat exposure to decrease or stabilize their emotional responses (42). This gradual emotional adaptation is reflected also by the perceived difficulty of the FH group, which declined over the course of the training despite the more challenging high-magnitude perturbations experienced by participants. In contrast, the LH group displayed a rise in their perceptions of training difficulty, as the gradually larger perturbations required greater balance corrections to be made.

## Limitations

The training perturbations were not completely unpredictable. Despite the variability in perturbation timing, participants were notified that each trial was starting, making it possible for anticipatory adjustments to be made before the perturbations. However, unpredictability in the perturbation direction meant that participants could not make direction-specific anticipatory adjustments. We assessed young, healthy adults, making our results not directly applicable to older adults or those with balance impairments. Future studies are needed to increase the generalizability of our results to those at greater risk of falling.

## Conclusions

This study showed that participants could improve balance recovery in response to perturbations in an untrained direction, regardless of intensity schedule. This suggests that RBT consisting of multi-directional perturbations may be more beneficial for generalizing improvements in reactive balance control than considerations regarding intensity schedule. Future research should focus on developing training protocols that incorporate a variety of perturbation directions.

