## Appendix 1 for "The impact of perturbation intensity schedule on improvements in reactive balance control in young adults: an experimental study"

**APPENDIX 1: Perceptions of perturbations questionnaire**

Please circle the corresponding number to indicate how strongly you agree or disagree with the statements below

| | Strongly disagree | | | Neutral | | | Strongly agree |
|---|---|---|---|---|---|---|---|
| I found the perturbation training stressful | 1 | 2 | 3 | 4 | 5 | 6 | 7 |
| It was easy to predict the next task | 1 | 2 | 3 | 4 | 5 | 6 | 7 |
| I enjoyed taking part in the training | 1 | 2 | 3 | 4 | 5 | 6 | 7 |
| The training was difficult | 1 | 2 | 3 | 4 | 5 | 6 | 7 |
| The perturbation tasks were always unexpected | 1 | 2 | 3 | 4 | 5 | 6 | 7 |
| I did not find the training challenging | 1 | 2 | 3 | 4 | 5 | 6 | 7 |
