## Appendix 2 for "The impact of perturbation intensity schedule on improvements in reactive balance control in young adults: an experimental study"

### APPENDIX 2: Analysis with participants who had protocol deviations removed

Five participants in the Low-high (LH) group, and 2 participants in the Variable (VA) group were removed from the analysis due to completing a different training protocol than intended (see details in the paper).

#### Arousal and perceptions of perturbations

There was a significant group-by-training block interaction effect for OMNI perceived exertion scores ($F_{4,27}$=7.50, p=0.0001). OMNI perceived exertion scores significantly declined between Training Block 1 and 2, and between Training Block 1 and 3 for the Fixed high (FH) group (Figure A2.1). There were no significant differences in OMNI perceived exertion scores between training blocks for the other groups, or between groups for any training block. These findings differ from analysis of the full sample, where there was a significant increase in OMNI perceived exertion scores between Training Blocks 1 and 2, and between Training Blocks 2 and 3 for the LH group.

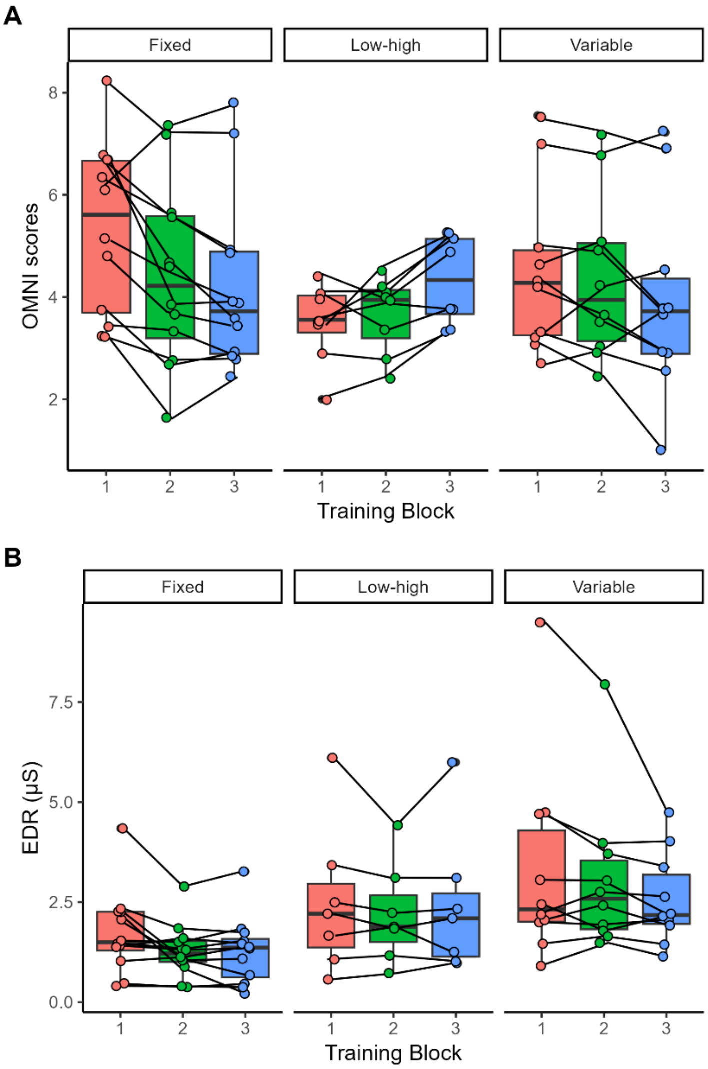


**Figure A1.2: Arousal and perception of perturbations during training.** OMNI perceived exertion scores (Panel A) and electrodermal responses (EDR, Panel B) for each group during the training blocks. Values plotted are individual data points for each participant, with line connecting data points for each participant. Box-plots show the group medians, and first and third quartiles.

There was no significant group-by-training block interaction effect ($F_{4,26}$=1.07, p=0.38) and no significant main effect of group for EDR ($F_{2,26}$=3.30, p=0.053). There was a significant effect of training block for EDR ($F_{2,26}$=4.33, p=0.018). Post-hoc testing revealed that EDR magnitude declined across all groups between Training Blocks 1 and 3 (Figure A2.1).

There were no significant differences between groups in perceptions of the perturbations at the end of the training session (Table A2.1).

**Table A2.1: Perceptions of perturbations.** Values presented are means with standard deviations in parentheses. The p-value is based on the comparison between groups using Kruskal-Wallis test.

| | FH | LH | VA | *p-value* |
|---|---|---|---|---|
| Stressful | 2.3 (1.3) | 3.3 (2.1) | 3.7 (2.4) | *0.38* |
| Easy to predict | 3.3 (1.8) | 2.6 (1.9) | 2.9 (2.2) | *0.72* |
| Enjoyment | 6.5 (0.9) | 6.4 (0.5) | 6.0 (1.1) | *0.36* |
| Difficulty | 1.9 (0.8) | 2.8 (1.5) | 3.3 (1.7) | *0.17* |
| Expected | 4.5 (1.2) | 4.6 (2.3) | 5.2 (1.4) | *0.57* |
| Did not find challenging | 5.3 (1.8) | 4.3 (1.9) | 3.9 (1.6) | *0.16* |

#### Primary outcome: step reactions

From ANCOVA, there was no significant between-group difference in number of steps ($F_{2,26}$=1.02, p=0.37) or proportion of multi-step reactions ($F_{2,26}$=0.01, p=0.99; Figure A2.2) post-training. Wilcoxon signed rank tests found that, overall, participants took fewer steps (Z=4.65, p<0.0001) and fewer multi-step reactions (Z=4.34, p<0.0001) post-training compared to pre-training.

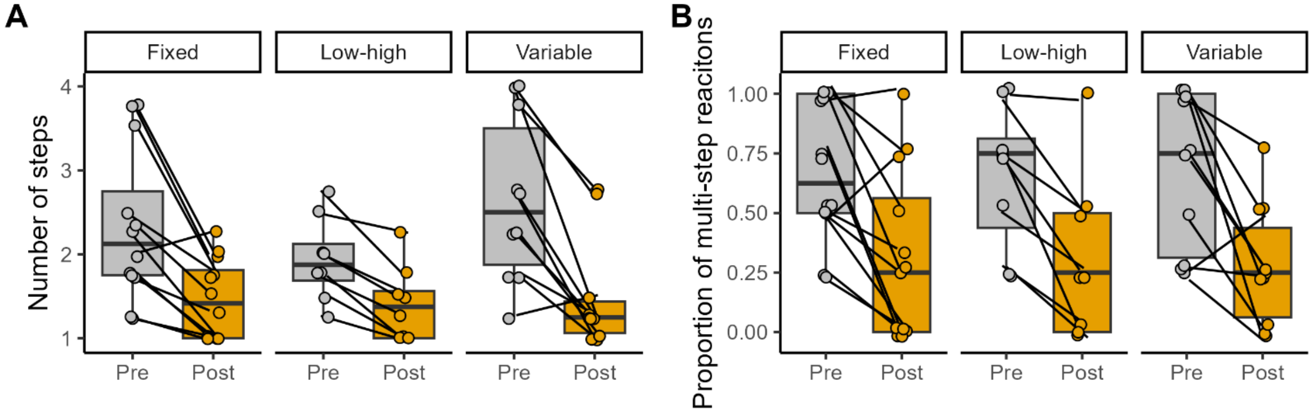


**Figure A2.2: Primary outcomes (step reactions).** Average number of steps (Panel A) and proportion of multi-step reactions (Panel B) for each group pre- and post-training. Values plotted are individual data points for each participant, with lines joining the pre- and post-training values for each participant. Box-plots show the group medians, and first and third quartiles.

When examining outcomes with the testing trial blocks, two way repeated measures ANOVAs found no statistically-significant group-by-trial interactions for average number of steps ($F_{6,27}$=0.87, p=0.52) or number of multi-step reactions ($F_{6,27}$=0.40, p=0.88), but there were significant main effects of trial order for both outcomes ($F_{3,27}$>2.77, p<0.047). Post-hoc testing revealed that average number of steps and proportion of multi-step reactions significantly declined between the 3rd pre-training trial and the 2nd post-training trial, and proportion of multi-step reactions was significantly lower for the 2nd post-training trial compared to the 4th pre-training trial. Average number of steps and proportion of multi-step reactions did not differ between the 3rd and 4th pre-training trials, or between the 4th pre-training trial and 1st post-training trial.

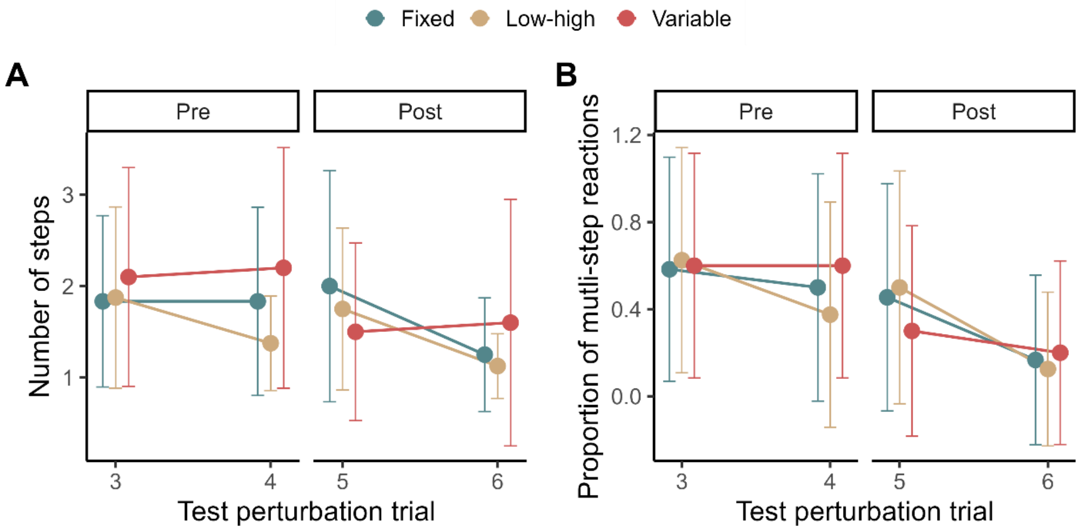


**Figure A2.3: Step reactions for individual trials.** Number of steps (Panel A) and proportion of multi-step reactions (Panel B) for the last 2 pre-training perturbations (trials 3 & 4), and the first 2 post-training trials (5 & 6). Values plotted are means with standard deviation error bars.

##### 3.2 Secondary outcomes

From ANCOVA, there were no significant between-group differences in any CoM-derived variables post-training (CoM displacement: $F_{2,26}$=0.28, p=0.76; AP-MoS: $F_{2,26}$=0.98, p=0.39; ML-MoS: $F_{2,26}$=0.72, p=0.50; Figure A2.4). There was a significant increase in ML-MoS pre- to post-training across all groups (Z=3.38, p=0.0004), but no significant change pre- to post-training for CoM displacement (Z=1.29, p=0.21) or AP-MoS (Z=0.42; p=0.69).

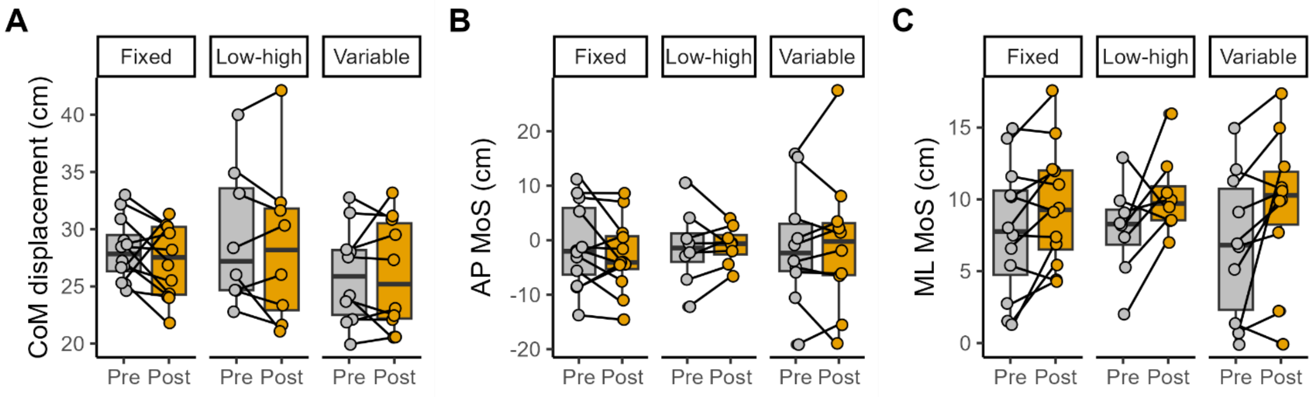


**Figure A2.4: Secondary outcomes (CoM-derived variables).** CoM displacement (Panel A), AP-MoS (Panel B), and ML-MoS (Panel C) for each group pre- and post-training. Values plotted are individual data points for each participant, with lines joining the pre- and post-training values for each participant. Box-plots show the group medians, and first and third quartiles.

There were no significant differences between groups in any spatio-temporal variables post-training (foot-off time: $F_{2,26}$=0.08, p=0.92; swing time: $F_{2,26}$=0.33, p=0.72; step length: $F_{2,26}$=0.42, p=0.66; step width: $F_{2,26}$=0.29, p=0.75; Figure A2.5). Likewise, there were no significant changes pre-to post-training for any spatio-temporal variables (foot-off time: Z=1.47, p=0.15; swing time: Z=1.14, p=0.26; step length: Z=0.35, p=0.73; step width: Z=1.51, p=0.13).

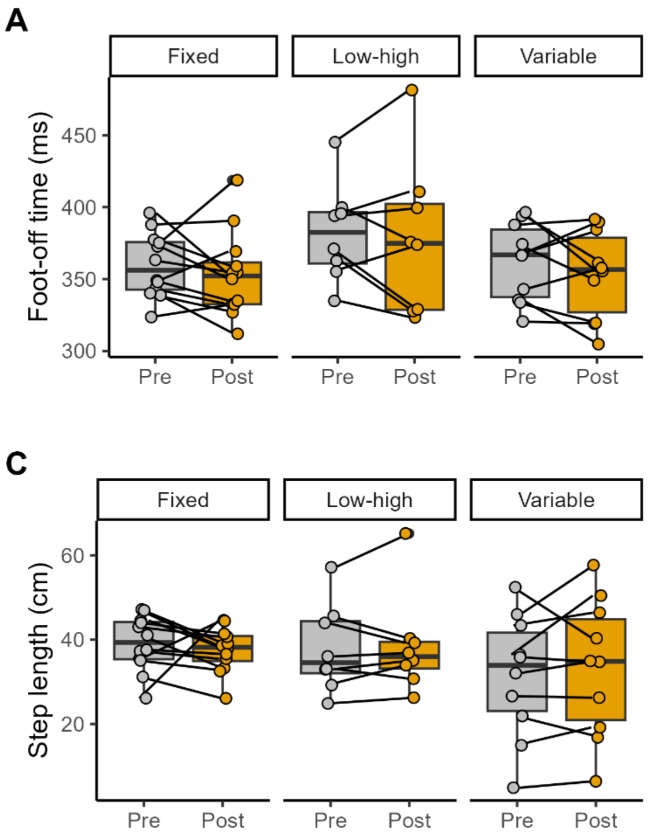


**Figure A2.5: Secondary outcomes (spatio-temporal variables).** Foot-off time (Panel A), swing time (Panel B), step length (Panel C), and step width (Panel D) for each group pre- and post-training. Values plotted are individual data points for each participant, with lines joining the pre- and post-training values for each participant. Box-plots show the group medians, and first and third quartiles.
